# Expression estimation and eQTL mapping for HLA genes with a personalized pipeline

**DOI:** 10.1101/365957

**Authors:** Vitor R.C. Aguiar, Jonatas E. Cesar, Olivier Delaneau, Emmanouil T. Dermitzakis, Diogo Meyer

## Abstract

The HLA (Human Leukocyte Antigens) genes are well-documented targets of balancing selection, and variation at these loci is associated with many disease phenotypes. Variation in expression levels also influences disease susceptibility and resistance, but little information exists about the regulation and population-level patterns of expression due to the difficulty in mapping short reads to these highly polymorphic loci, and in accounting for the existence of several paralogues. We developed a computational pipeline to accurately estimate expression for HLA genes based on RNA-seq, improving both locus-level and allele-level estimates. First, reads are aligned to all known HLA sequences in order to infer HLA genotypes, then quantification of expression is carried out using a personalized index. We use simulations to show that expression estimates are not biased due to divergence from the reference genome. We applied our pipeline to GEUVADIS dataset, and compared the quantifications to those obtained with reference transcriptome, and found that a substantial portion of the variation captured by the HLA-personalized index in not captured by the standard index (23%). We describe the impact of the HLA-personalized approach on downstream analyses for seven HLA loci (*HLA-A, HLA-B, HLA-C, HLA-DPB1, HLA-DQA1, HLA-DQB1, HLA-DRB1*). Although the influence of the HLA-personalized approach is modest for eQTL mapping, the p-values and the causality of the eQTLs obtained are better than when the reference transcriptome is used. Finally, we integrate information on HLA-allele level expression with the eQTL findings to show that the HLA allele is an important layer of variation to understand HLA regulation.

## Introduction

The HLA region is the most polymorphic in the genome, and also shows the greatest number of disease associations, which has made it very well characterized at the genomic, population and functional levels [1, 2]. Decades of research have also shown that the HLA genes are targets of natural selection, likely a consequence of their role in responding to pathogens [2, 3]. This combination of evolutionary and biomedical interest has resulted in an extensive catalogue of HLA variation in human populations, with the identity and population frequency of HLA alleles defined for various populations [4, 5, 6, 7].

Variation within or near HLA genes has been convincingly linked to disease phenotypes, providing an extensive list of SNPs within or near HLA genes associated with resistance and susceptibility to both autoimmune and infectious diseases [2, 8, 9]. Although in many cases the mechanistic basis of the associations remain poorly understood, there has been an effort to identify whether associations can be linked to features such as variation at the level of specific amino-acids, HLA alleles, HLA haplotypes, non-coding variants near HLA genes, or with HLA expression levels (reviewed in [2, 9, 10]).

In some instances, the association of one feature results from the fact that it tags another feature. For example, [11] showed that HLA expression levels influence Parkinson disease, and that classical associations with HLA alleles/haplotypes are sometimes driven by eQTL SNPs. [12] and [13] showed that HLA-C surface expression levels correlate with HIV load, and [14] observed an association of *HLA-DPB1* with HBV infection. In both cases, some HLA alleles may have a protective effect because of their relative contribution to the overall gene expression. Other studies showed that a combination of two features may be important. For example, [15] observed an association of mortality of transplant recipients with increased expression of *HLA-C* alleles with certain amino-acid residues.

An understanding of how HLA expression varies among individuals, and the identification of genetic variants involved in the regulation of expression, will play a central role in understanding the contribution of HLA genes to normal and disease phenotypes. However, until recently, little information existed about the regulatory variation and population-level expression patterns of HLA genes, a result of the difficulty in quantifying expression for genes which show an unusually high polymorphism and are members of a multi-gene family [8, 16].

Efforts have been made to develop antibody-based methods to quantify HLA protein on the cell surface [13, 17], or hybridization-based approaches to quantify mRNA, such as qPCR [18, 19] and microarray methods [20]. However, the optimization of PCR reactions for a large set of allele-specific primers, and the design of probes that span the diversity of possible variants represent a technically challenging and labor-intensive undertaking. In addition, qPCR technologies are not appropriate for comparison of expression levels among different loci, an important concern when seeking to understand how expression of HLA genes responds to environmental challenges.

Despite the difficulties with these approaches, the results obtained to date indicate the importance of understanding the genetic architecture underlying HLA expression [12, 13, 15, 21, 22, 23]. However, they do not take advantage of the large amount of RNA-seq data generated by studies of whole transcriptomes in large samples [24, 25, 26], often involving populations from various regions of the world, which represents an attractive resource to investigate HLA expression.

Although such whole-transcriptome RNA-seq studies do provide expression estimates for HLA genes, they bring new challenges. RNA-seq pipelines may provide biased expression estimates for two reasons: 1) many short reads originating from genes with extreme polymorphism fail to map to the reference genome, due to high degree of variation (which results in a large number of mismatches between the reference genome and that of most individuals being analyzed), and 2) the presence of paralogues makes it difficult to map a read uniquely to a specific gene, leading to the exclusion of many reads. This raises concerns about the utility of RNA-seq approaches to quantify HLA expression, given that these loci represent both the extreme of polymorphism in the human genome and are part of a multi-gene family [27, 28, 29].

A strategy to overcome these challenges is the mapping of reads to an HLA-personalized reference, rather than to a single reference genome. For example, seq2HLA is a tool developed by [30] to provide *in-silico* HLA types and expression estimates, and later applied to demonstrate that different tumors types are associated with different HLA expression levels [31], and also to provide a large catalog of HLA expression in 56 human tissues and cell types [32]. `AltHapAlignR` [33] is another recently described algorithm which infers the MHC references which are the closest to the individual’s MHC haplotypes, and maps reads to them. The authors reanalyzed the GEU-VADIS dataset [24], and provided comparisons with conventional read mapping, showing an improvement in accuracy with the HLA-tailored pipeline.

In this article, we compare the results of conventional and HLA-personalized pipelines to analyze the reliability of RNA-seq quantification. We discuss for the first time the impact of accurate estimation of HLA expression on downstream analyses such as eQTL mapping and allele-specific expression. We show that it is possible to adapt different computationally efficient methods available to work under the strategy of a personalized reference to reliably quantify HLA expression from RNA-seq data. We find that implementations with either a conventional read mapper [34] or a pseudoaligner [35] show similar expression estimates.

We use simulations to assess accuracy, showing that HLA-personalized pipelines are more accurate than conventional mapping, and apply the tool to reanalyze RNA-seq data of the GEUVADIS dataset of Lymphoblastoid Cell Lines (LCLs) [24]. We then sought to evaluate the impact of more accurate expression estimates on downstream analyses by carrying out a detailed survey for allele-specific expression and eQTL mapping at 7 classical HLA loci *(HLA-A, HLA-B, HLA-C, HLA-DPB1, HLA-DQA1, HLA-DQB1*, and *HLA-DRB1*).

Surprisingly, we find that conventional RNA-seq pipelines provide gene-level expression estimates and identify eQTLs which are highly correlated with those obtained under the HLA-personalized approach. However, we identify gains of using a pipeline tailored for HLA diversity. These include more accurate expression estimates, eQTLs with higher probabilities of being causal, and expression estimates at the HLA allele level.

## Results

### HLA expression quantification from RNA-seq data

We developed the `HLApers` pipeline (for HLA expression with personalized genotype) to measure HLA expression from whole-transcriptome RNA-seq data. The pipeline can use either (1) a suffix array-based read mapper (STAR [34]) followed by quantification with Salmon [36] (henceforth called STAR-Salmon), or (2) a pseudoaligner with built-in quantification protocol (kallisto [35]). The key feature of our implementation is the use of an index supplemented with a set of sequences covering the breadth of known HLA sequences (see Materials and methods). We implemented a two-step quantification approach, where (1) we identify the HLA alleles that maximize the read counts at each locus allowing us to infer the genotype which is present *(in-silico* genotyping), and (2) we use this HLA genotype to create a personalized index which we use to quantify expression (Figure 1).

**Figure 1:**
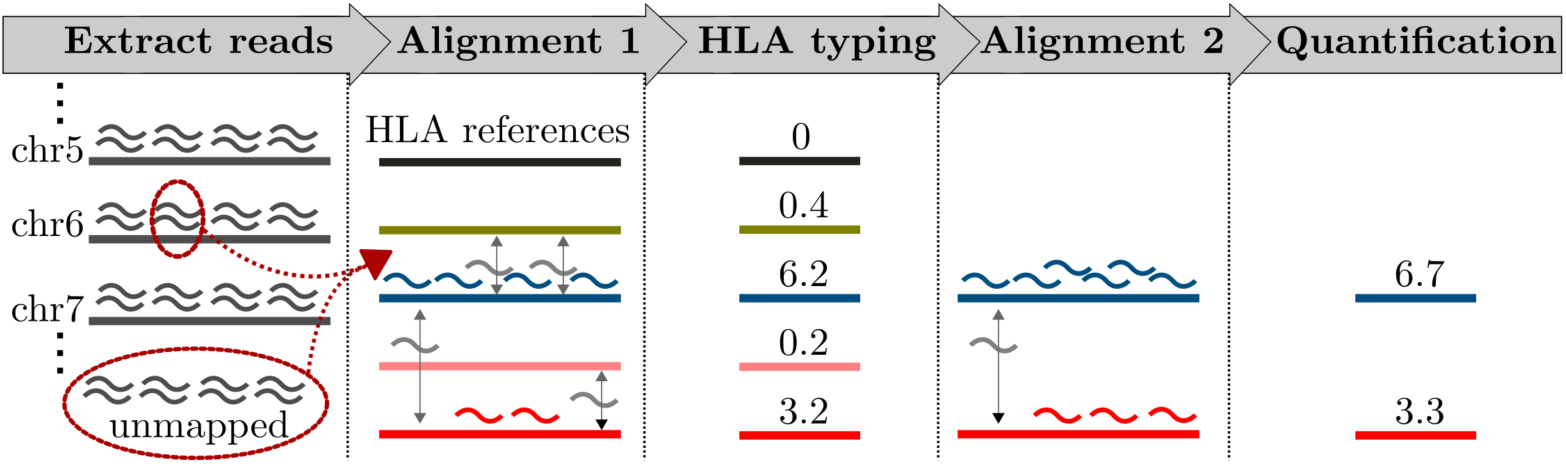
Schematic representation of the HLApers pipeline to estimate HLA expression. The reads generated by RNA-seq are represented by curved shapes. The pipeline can use as input either reads in fastq format or extracted from a BAM file *(Extract reads*). These reads are aligned to an index containing all known HLA alleles *(Alignment 1*), and we pick the alleles for which most reads align to infer the HLA genotypes of the individual at that locus, in this example a heterozygote carrying a blue and red allele *(HLA typing*). In a second step a new alignment is performed, using as an index only the alleles inferred to be present in that individual at this locus *(Alignment 2*). Expression is estimated based on the number of reads aligning to each allele, using a statistical model that accounts for instances where reads align to multiple HLA alleles or genes *(Quantification*).

For both first and second steps of our pipeline a read can align to more than one allele or gene (due to their sequence level similarity). Instead of discarding such reads (as in [33]) or evenly splitting them among the compatible references (as in [30]), we use maximum likelihood estimates of expression obtained by an expectation-maximization (EM) algorithm, which is implemented within Salmon and kallisto. This procedure probabilistically assigns reads to each reference in the index, in a way that accounts for reads that align to more than one gene or allele [35, 36]. The pipeline is available at https://github.com/genevol-usp/HLApers.

Because the *in-silico* typing is an important step for accurate expression estimates, we assessed the concordance between our RNA-seq based HLA typing and the HLA allele calls experimentally determined by [6] using Sanger sequencing. The concordance was higher than 97% for all of the 5 HLA genes compared (Table S1). This is consistent with previous results showing that RNA-seq provides reliable HLA alleles calls [30, 37, 38, 39, 40].

### Expression quantification for simulated data

We next investigated how using the HLA-personalized index affects the quantifications of HLA expression. To this end we simulated an RNA-seq experiment using the Polyester package [41], with a dataset of 50 individuals and read counts matching those of the observed data, with the same read length as in the original data (paired-end 75bp reads) and without bias.

We analyzed the simulated dataset using three different methodologies: (1) Using the two-step approach in HLApers, first we inferred the personalized HLA genotype and then aligned reads to it, (2) Alignment to the reference transcriptome (Gencode release 25; primary assembly), and (3) Alignment to the reference genome (GRCh38). In approaches (1) and (2), the alignment is followed by expression quantification using Maximum Likelihood (ML), which provides a statistical framework for dealing with multimap reads, whereas in approach (3) quantification is performed using only uniquely mapped reads.

For each HLA locus and methodology, we assessed the proportion of simulated reads which successfully aligned. Because previous studies for genome sequencing identified a correlation between mapping success and the number of mismatches between the HLA allele an individual carries and the reference genome [28], we analyzed how alignment success behaves as a function of the number of mismatches between each HLA allele and the reference genome (Figure 2).

**Figure 2:**
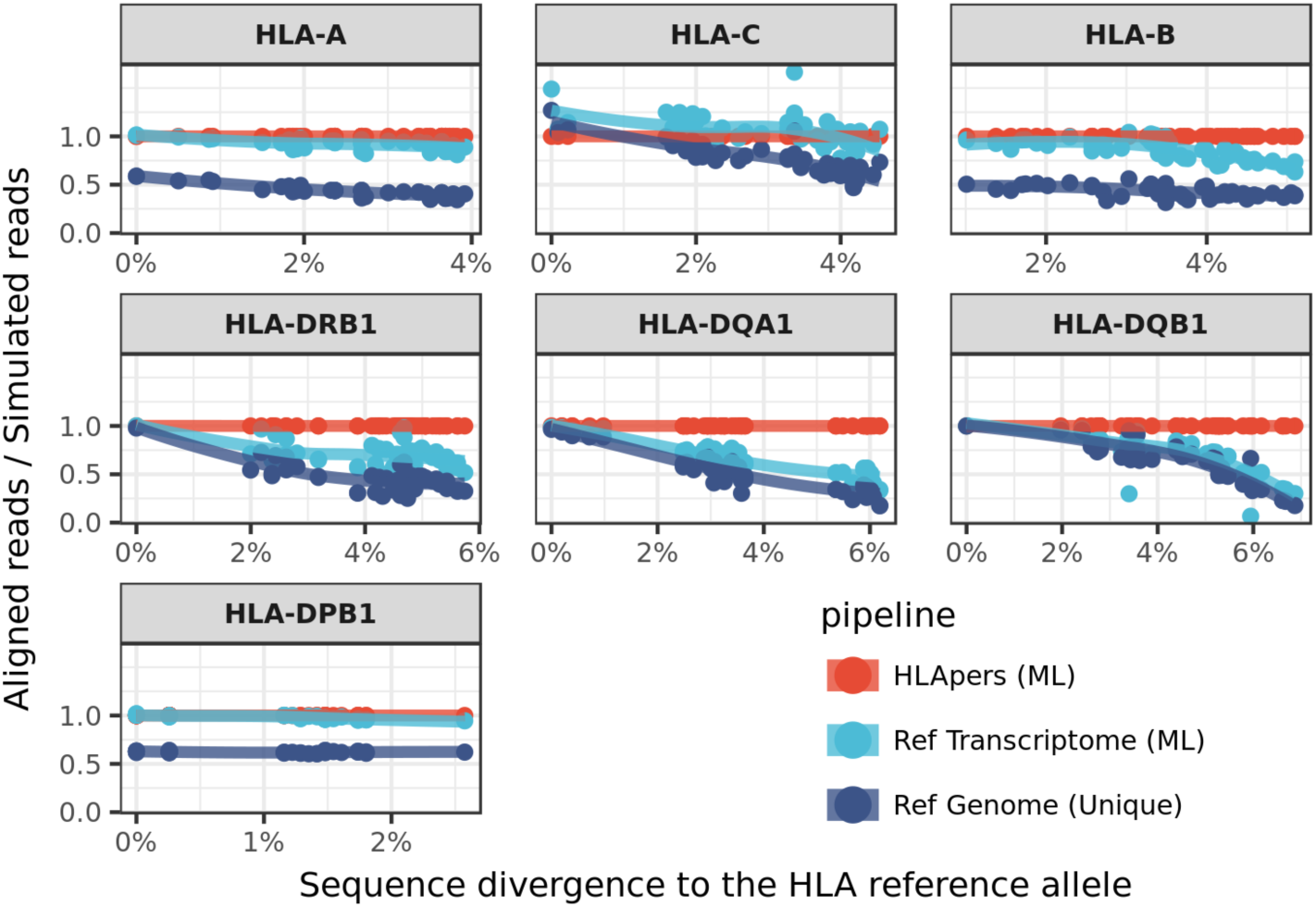
Alignment success with three different methodologies. The proportion of simulated reads which successfully aligned is defined for each locus by the ratio of the estimated read counts to the number of simulated reads (y-axis). These values are displayed as a function of the divergence of the HLA allele to the reference genome (x-axis). Simulated reads were processed through 3 different pipelines: (1) alignment to the personalized HLA sequences *(HLApers (ML))*, (2) alignment to the reference transcriptome *(Ref Transcriptome (ML*)), (3) alignment to the reference genome considering only uniquely mapped reads *(Ref Genome (Unique*)).

The use of an HLA-personalized index results in the largest proportion of successfully aligned reads, no matter how different the allele carried by the individual is from the allele in the reference genome. This is expected, since the personalized HLA component guarantees that a sequence close or identical to that originating the read will be present.

When the alignment was performed using the reference transcriptome, there was a marked reduction in the proportion of successfully aligned reads for *HLA-B, HLA-DRB1, HLA-DQA1, HLA-DQB1*. In all cases, alignment success decreased for alleles which showed progressively greater proportions of mismatches with respect to the reference genome.

Finally, when using uniquely mapped reads there was a massive read loss regardless of the divergence for *HLA-A, HLA-B* and *HLA-DPB1*, and a lower proportion of successfully aligned reads across all surveyed loci. This shows that both discarding multipmaps, as well as not including a personalized index, have a negative impact on mapping success.

### Analysis of the GEUVADIS dataset

Having demonstrated that the presence of an individual’s HLA alleles in the index has a substantial impact on the success of read alignment with simulated data (Figure 2), we set out to address two questions with real data. First, we examined how expression varies among HLA loci, when the personalized index is used (Figure 3). Secondly, we compared expression estimates with and without the use of the personalized index, so as to evaluate the impact of its usage on real data (Figure 4).

**Figure 3:**
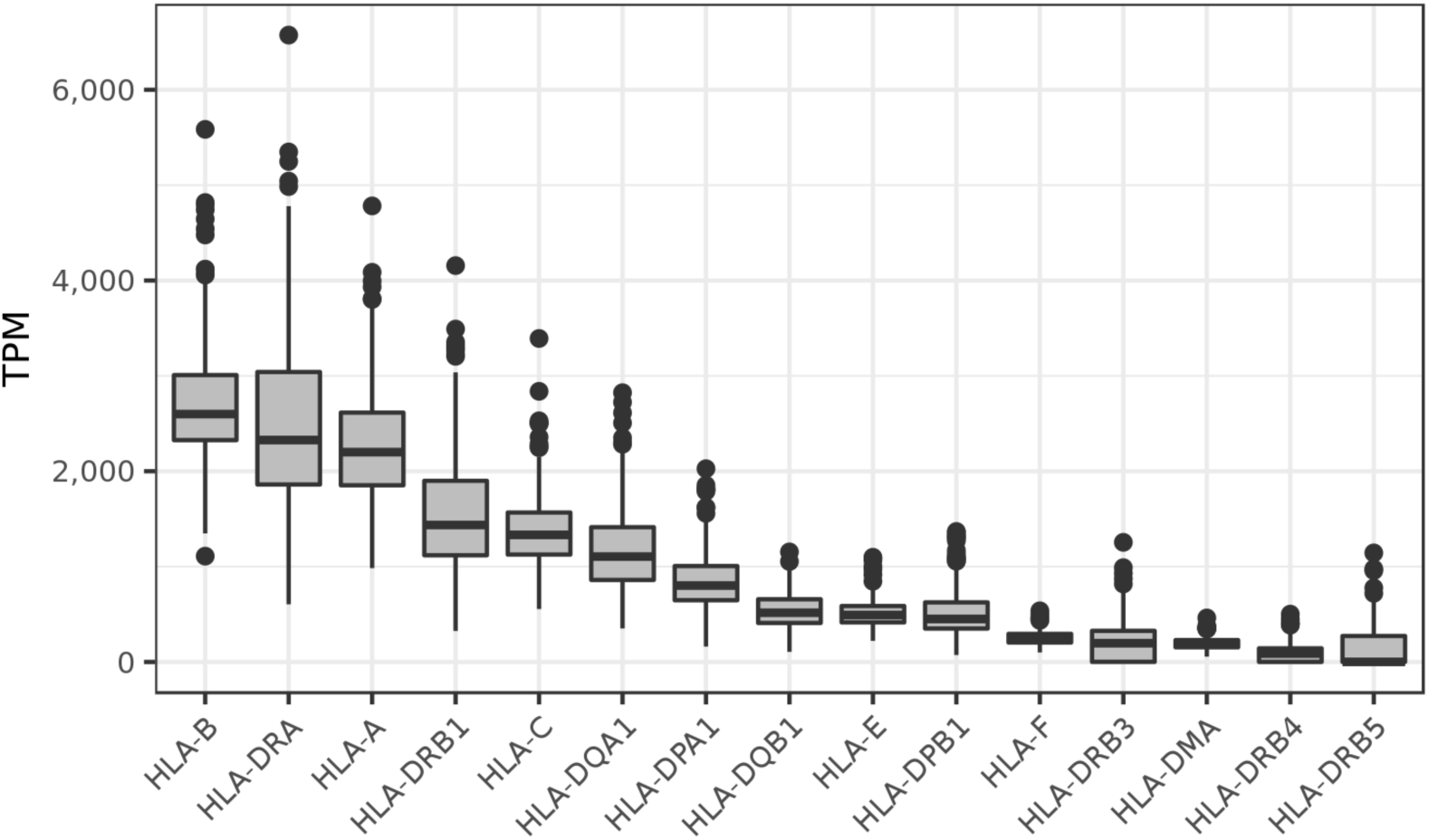
Gene-level expression of 358 European individuals in the GEUVADIS dataset [24]. Here we show all HLA genes with expression levels ≥ 100 TPM. TPM: Transcripts per Million.

**Figure 4:**
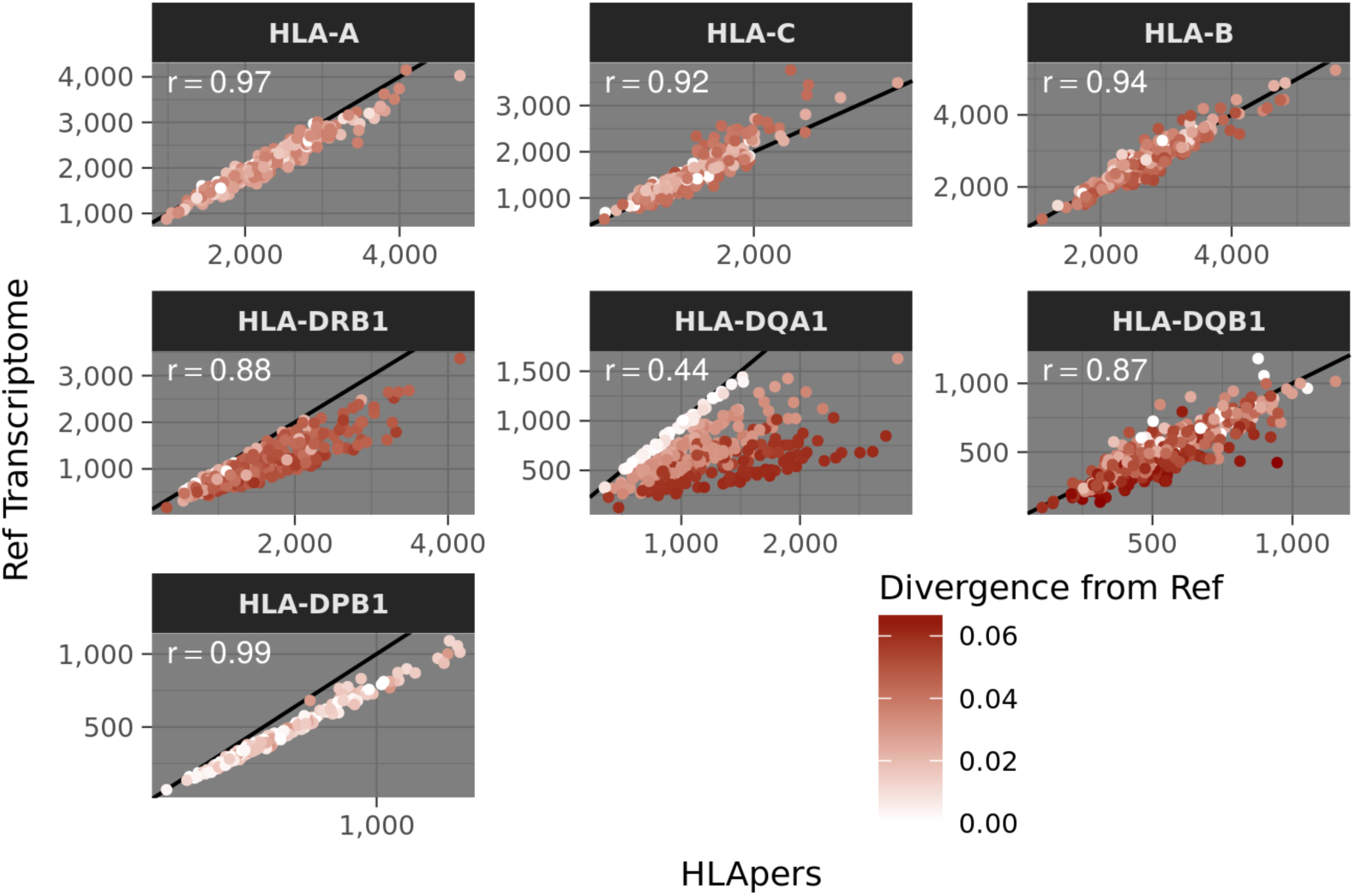
Expression estimates using personalized HLA genotypes (x-axis) versus the reference transcriptome (y-axis) in the index. For each individual we identify the average divergence (proportion of mismatches) with respect to the reference genome, indicated by the red gradient. For all loci with low correlations between expression quantification methods, there is a marked drop in expression estimates using the reference transcriptome when the alleles are most divergent from the reference. Expression estimates are in TPM (Transcripts per Million).

By summing the estimates for the 2 alleles at each HLA locus, we obtain gene-level expression estimates (Figure 3). We observe that *HLA-B* is the highest expressed gene overall. Among the Class I genes, *HLA-B* is followed by *HLA-A* with similar levels, and by *HLA-C* which has about 50% of the expression levels of *HLA-B*. For Class II genes, *HLA-DRA* is the most highly expressed. Although we observe a general concordance with the original GEUVADIS quantifications [24], for some loci the ordering and fold change among genes are different. For example, for the GEUVADIS original quantifications, *HLA-B* is twice as expressed as *HLA-A*, and *HLA-DPA1* is more expressed than *HLA-DRB1* (Figure S1). Our results are more in accordance with previous HLA-personalized approaches which shows, for example, that HLA-DR is always more expressed than HLA-DP and HLA-DQ [32].

We found that, although the expression estimates using the reference transcriptome were usually lower than using the personalized index, the correlation between indices was greater than 0.87 for every locus except for *HLA-DQA1* (Figure 4). However, a substantial portion of the variation captured by the HLA-personalized index is not captured by the reference transcriptome index (average 1 − *R*^2^ = 0.23). The loci with the lowest correlations between indices (*HLA-DQA1*, *HLA-DQB1* and *HLA-DRB1*), are also those with the greatest read loss when divergence from the reference allele is high, consistent with the pattern observed in the simulation (Figure 2).

We then investigated if the bioinformatic tool used to quantify expression influences expression estimates. When comparing estimates obtained using the personalized index, those from STAR-Salmon and those from kallisto had an average correlation of *r* > 0.99 for read counts, dropping to *r* = 0.9 for TPM estimates, likely due to different bias correction models (Figure S2). Overall, these results show that the key features influencing alignment success are the use of a personalized index and the statistical treatment of multimaps (as opposed to discarding them), with the specific alignment tool being less influential.

### eQTL analysis of HLA loci

The key role played by HLA loci in the immune response and their strong and abundant associations with infectious and autoimmune diseases have motivated studies to uncover their regulatory architecture.

Here, we use our approach based on a personalized index to obtain accurate expression estimates, together with genotype data from the 1000 Genomes Project [42], to identify SNPs which are associated with variation in expression levels (eQTLs).

Because multiple SNPs can affect expression, it is interesting to identify independent contributions made by distinct SNPs. This has previously been done by using a best eQTL (i.e., the one with the most extreme p-value) as a covariate in subsequent searches for an additional variant. Here, we use a conditional analysis available in `QTLtools` [43] in order to identify groups (or “ranks”) of SNPs associated with independent signals. For each rank, we identified the site with the most extreme association (Figure 5).

**Figure 5:**
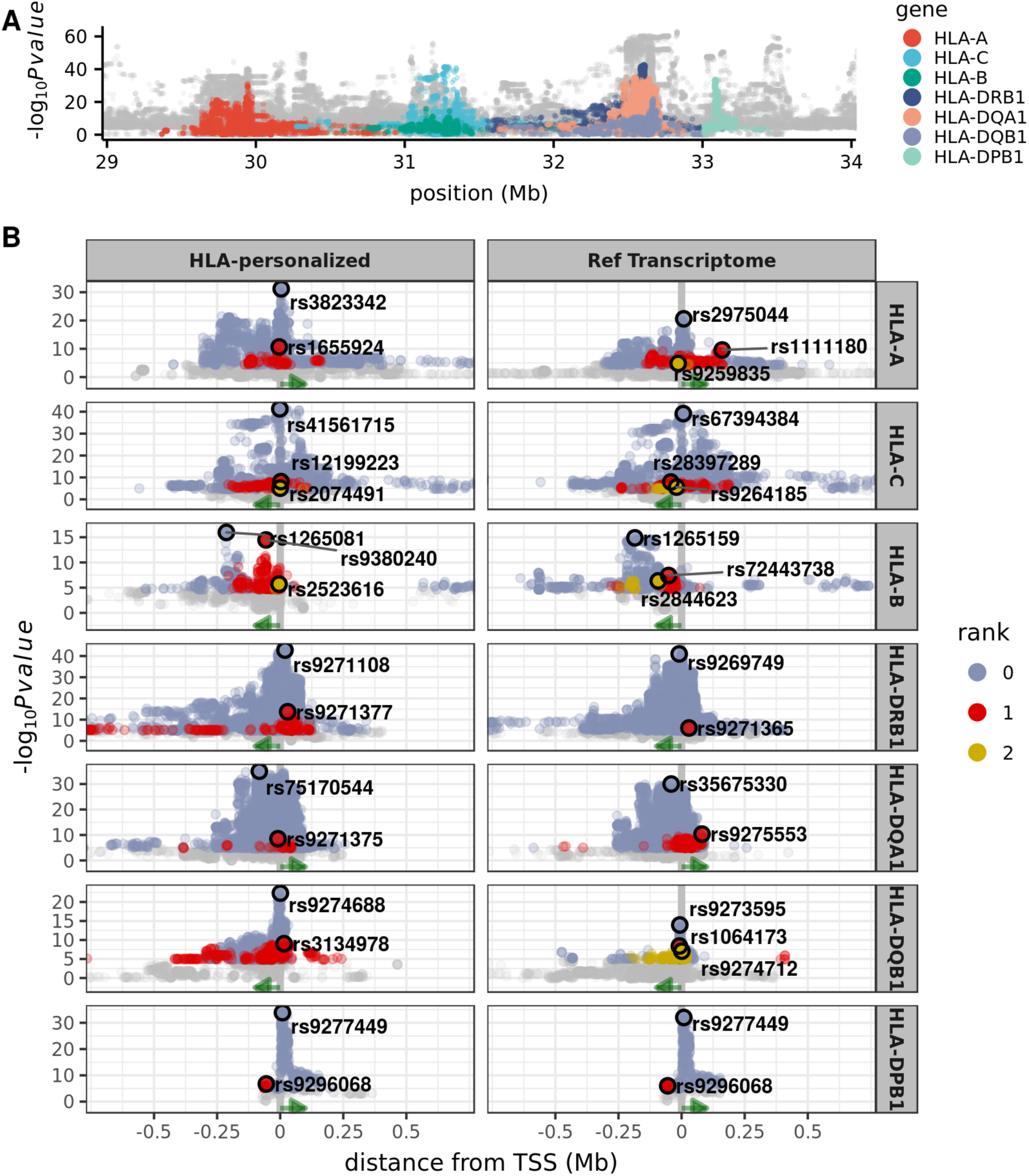
eQTLs for HLA loci. **A** Distribution of p-values for the association between genotypic and expression variation along the MHC region colored by gene. **B** Distribution of p-values around the transcription start site (vertical line at x = 0), for eQTLs mapped with the HLA-personalized or reference transcriptome pipelines. Points are colored according to the rank of association (with gray for non-significant associations at an FDR of 5%). The best eQTL of each rank is circled in black. Green arrows indicate the orientation of transcription.

#### Differences across indices

Although expression estimates obtained using HLA-personalized and reference transcriptome indices are significantly correlated, there is a global reduction in expression estimates based on the reference transcriptome (Figures 2 and 4). We therefore examined if these differences in expression across indices have a direct effect on the eQTLs identified.

Even if eQTLs across studies or pipelines are different, they may both tag the same biological signal (possibly driven by the same causal site). Therefore, a simple comparison of eQTL sharing may be inappropriate. We thus used a Regulatory Trait Concordance (RTC) analysis [43, 44] to test if eQTLs from different pipelines tag the same signal (indicated by RTC score ≥ 0.95).

We observed that the lead eQTLs capture the same biological signals across indices for all loci except *HLA-DRB1* (Figure 5 and Table S2). Despite this sharing of eQTL discoveries, the p-values for the eQTLs mapped under the HLA-personalized pipeline are in almost all cases more extreme than those from the reference transcriptome (Figure 5).

#### eQTL sharing with previous studies

Next we investigated whether the eQTLs for HLA genes which we mapped share the same causal signal as those reported by previous studies. We queried three large studies of LCLs [24, 26, 45], including the original GEUVADIS publication, and also HLA-targeted studies [12, 17, 21, 46].

Although the variants which we mapped as eQTLs were not reported among the top associations in previous studies, we find that they likely share the same signal as previous lead eQTLs for Class I genes and *HLA-DPB1* (Table S3). However, for *HLA-DQA1, HLA-DQB1*, and *HLA-DRB1*, our eQTLs do not share the same signal as previously reported eQTLs. This is consistent with our finding that for these genes the expression estimates are affected by divergence from the reference genome (Figures 2 and 4), which may have led to incorrect estimation of expression in previous studies.

However, as we discuss below, the lead eQTLs mapped with the HLA-personalized pipeline have low causal probabilities (for *HLA-DQA1* and *HLA-DRB1*, Figure S4), or no functional annotation from ENCODE (for *HLA-DQA1*, Table S4), which shows a general difficulty in mapping eQTLs for these genes in the low/moderate sample sizes.

#### Putative function of eQTLs

To gain insight into possible function of the eQTLs mapped with the HLA-personalized pipeline, we examined their location with respect to the associated HLA gene and functional annotation for LCLs in ENCODE [47], such as transcription factor binding sites (TFBS), DNase I hypersensivity sites (DHSs), and chromatin modifications.

The lead eQTL was located no further than 20kb from the gene for *HLA-A*, *HLA-C, HLA-DRB1, HLA-DQB1*, and *HLA-DPB1*, whereas for *HLA-DQA1* it was 82kb upstream the transcription start site (TSS), but within a cloud of associations which extends to the TSS, and for *HLA-B* the lead eQTL was located 210kb away, an unusually large distance with respect to that of the other loci, but still within a range found in the genomewide eQTL data (where 11.1% of the eQTLs lie at that distance or further).

Considering all ranks of association, we mapped 16 eQTLs in total, of which 5 are part of TFBS/DHSs. Regarding chromatin modification, 10 eQTLs were located within regions with marks of active promoters, enhancers or other distal elements, transcribed genes, or elements with dynamic chromatin (Table S4).

We also used RTC to evaluate if eQTLs identified in our study capture the same signals as CRD-QTLs identified by [45]. CRDs (for Cis-Regulatory Domains) are segments of the genome defined by chromatin activity and formed by coordination between nearby regulatory elements [45] (See Figure S3 for the location of HLA genes in respect to CRDs in the region). A high RTC score between a CRD-QTL and an eQTL indicates a shared influence on the formation of a CRD and gene expression. Overall, 6 out of 16 eQTLs we identified mark the same signal as CRD-QTLs. Interestingly, the lead eQTL for *HLA-B*, which is 210kb away from the gene, has an RTC score of 0.99 with a variant associated with the activity of the CRD linked to *HLA-B* (Table S5).

Studies have revealed a significant overlap of eQTLs and GWAS variants, which provides insight into a biological basis for the GWAS association [10, 24, 48, 49, 50]. Most GWAS variants are non-coding, and in those cases it is difficult to know which variant is causal and which gene is modulated, complicating the identification of the direct effects of variants on disease etiology [44, 51]. Mapping of eQTLs, by identifying variants involved in modulating expression, offers an additional layer of information for interpreting GWAS hits. Thus, we sought to investigate the overlap between eQTLs we mapped and GWAS signals.

We found that nearly all of our eQTLs, according to the RTC score, likely share the same biological signal as GWAS hits (Table S6), which shows an extensive implication of regulation of gene expression in disease phenotypes at the HLA region.

#### Causality

A key challenge for eQTL studies is identifying which variants are causally related to variation in expression, and which are simply genetically correlated with the functional variant. One approach to identify causal variants is to use empirical and simulated data on eQTLs in a probabilistic framework, to estimate the probability that a variant is causal (implemented in CaVEMaN [49]). We used this method to estimate the probability that the eQTLs we mapped are causal, and to evaluate if more accurate expression estimates could lead to improved mapping of causal variants.

For 10 out of 14 eQTLs (excluding *HLA-DPB1*, for which eQTLs are the same across indices), we found that the HLA-personalized pipeline produces eQTLs with higher probabilities than the reference pipeline. However, most variants do not reach probabilities of 0.8, indicating that even though the personalized pipeline produces more accurate expression estimates, larger samples sizes are important to improve fine mapping (Figure S4).

### HLA allele-level analysis

Transcriptome studies can quantify expression for various biological features: individual SNPs, exons, isoforms, genes. In the case of HLA loci, a natural unit of interest is the HLA allele. Our HLA-personalized pipeline provides expression estimates for individual alleles, since it is the allelic sequences which are included in the index. The immunogenetics literature has shown that many HLA alleles are associated with specific phenotypes of evolutionary and medical importance [reviewed in 2, 9, 10]. Gauging information about the expression levels of alleles can therefore provide an additional layer of information.

#### HLA allele lineages and eQTLs

To investigate HLA regulation at the allele level, we estimated expression levels for HLA alleles, and we inferred haplotypes that spanned HLA loci and eQTLs, so that the phasing between these was known.

We then grouped alleles in “lineages”, which comprise groups of alleles which are evolutionarily and functionally related, since the large number of alleles would make sample sizes per allele too sparse. Although the expression of individual allelic lineages is highly variable among individuals, there is an overall significant difference in expression among lineages (Welch’s ANOVA p-values ranging from 3.7×10^−8^ for *HLA-DPB1* to 6×10^−51^ for *HLA-DQA1*).

We investigated the degree to which the mapped eQTLs account for variation in expression levels of HLA alleles. Frequently, we observe that more highly expressed HLA alleles are on haplotypes carrying the eQTL allele associated with increased gene-level expression (red dots in Figure 6). At *HLA-C*, the rs41561715-T allele is exclusively associated with the C*04 lineage. This variant is located within the gene, and in fact there are 14 variants tied in p-value marking the C*04 lineage, distributed from upstream to downstream the gene.

**Figure 6:**
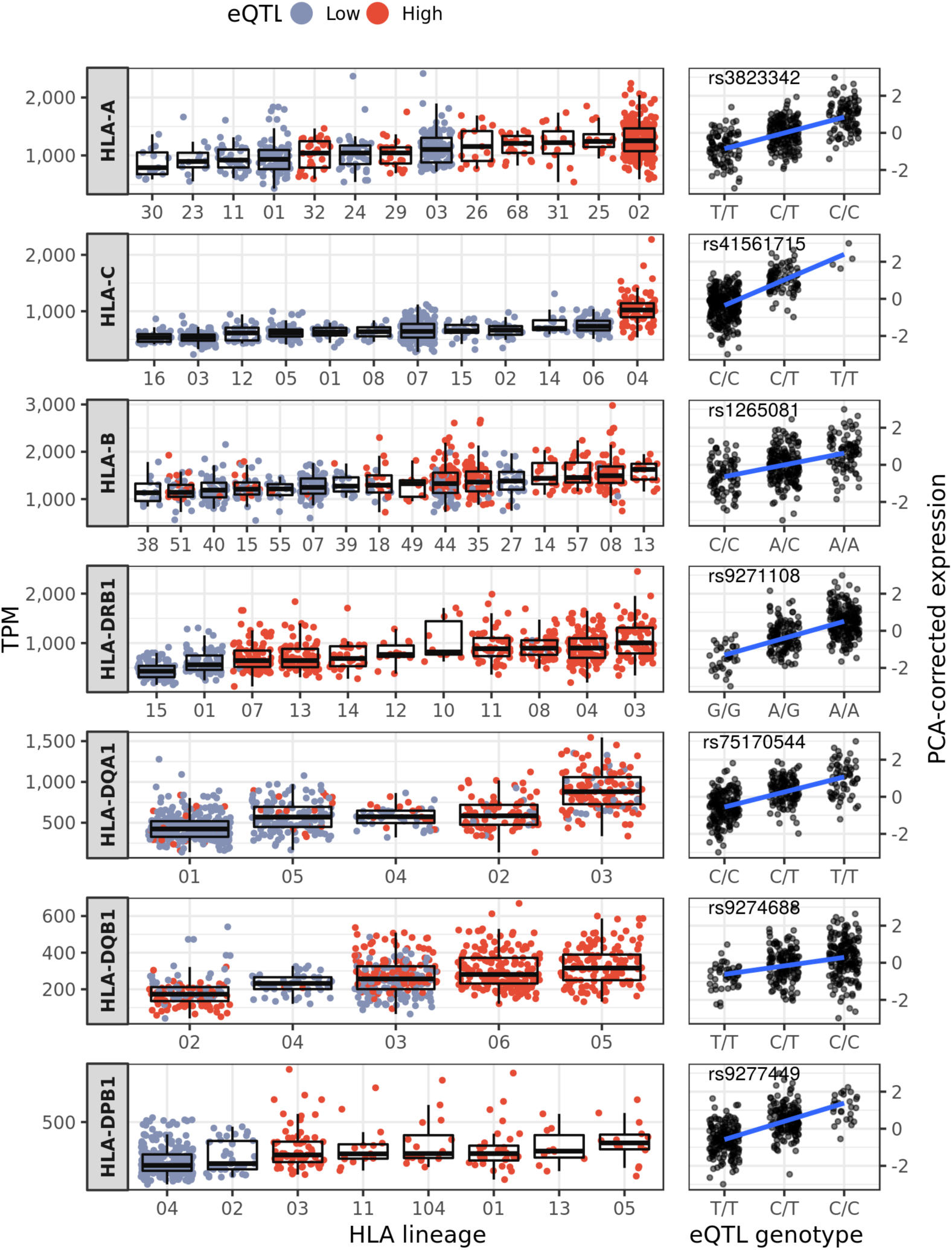
Expression levels for different HLA lineages (left-hand side panels), and eQTL genotypes (right-hand side panels).

Another instance is seen at *HLA-DPB1*. The alleles at this locus are divided into two clades, marked by the SNP variant rs9277534 in the 3’-UTR [23, 52]. The alleles DPB1*04 and DPB1*02 are on the same haplotype as rs9277534-A, and were associated with low expression, whereas the other alleles are on haplotypes carrying rs9277534-G, were associated with higher expression [23, 52], and may increase the risk of persistent Hepatitis B virus (HBV) infection [14]. Here we mapped rs9277449 as the lead eQTL for *HLA-DPB1*. This variant is located 2kb away and has a *D*′ = 1 with rs9277534, and although the RTC analysis for shared causal signal was not significant (RTC = 0.89), and rs9277449 is likely not the causal SNP (Figure S4), these two variants are non-independent (both are contained among the rank 0 variants (blue) in Figure 5) and, strikingly, rs9277449 also separates the two clades of alleles based on expression (Figure 6).

These results show that reliable eQTL mapping may help explain differences in expression among HLA alleles, and help identify genetic variants which contribute to disease risk.

The expression levels for HLA lineages which we report are different from those of previous studies, showing a distinct ranking of lineages by expression levels. This may be explained by different techniques (RNA-seq vs. qPCR), the samples used, or the fact that imputation of expression was used for heterozygous genotypes in previous studies [13, 18]. Even in a comparison with [33], which also reanalyzed the GEUVADIS dataset with an HLA-personalized approach, we observe only a moderate concordance in the ordering of lineages by expression level. This is likely due to several differences between our approach and theirs, including the strategy for alignment ([33] use the 8 MHC region haplotypes to guide quantification, while we directly use the entire known HLA diversity for this task), and the treatment of reads mapping to multiple loci or alleles ([33] discard these reads, while we use a maximum likelihood approach to measure their contribution).

#### Allelic imbalance

There is increasing interest in understanding if gene expression is unbalanced (i.e., one allele is more expressed than the other), and if so, how this contributes to interpretations of disease phenotypes and natural selection [53]. In the case of HLA genes, for which heterozygote advantage is a selective regime with strong theoretical support, extremely unbalanced expression poses a theoretical challenge, since being heterozyote would not be advantageous if only one allele is expressed. We have used our expression estimates to quantify allelic imbalance for HLA genes.

Using the expression data at the HLA-allele level we found a low level of asymmetry in expression, with 77% of the heterozygotes having ASE between 0.4 and 0.5, with Class II genes showing a distribution with a larger variance (Allele-specific expression (ASE) was defined as the proportion of the gene expression attributed to the less expressed HLA allele, Figure 7), and we see no instances of extreme imbalance as recently reported for SNPs in HLA genes [54]. Therefore, the extreme imbalance seen at the SNP level does not hold at the HLA allele resolution.

**Figure 7:**
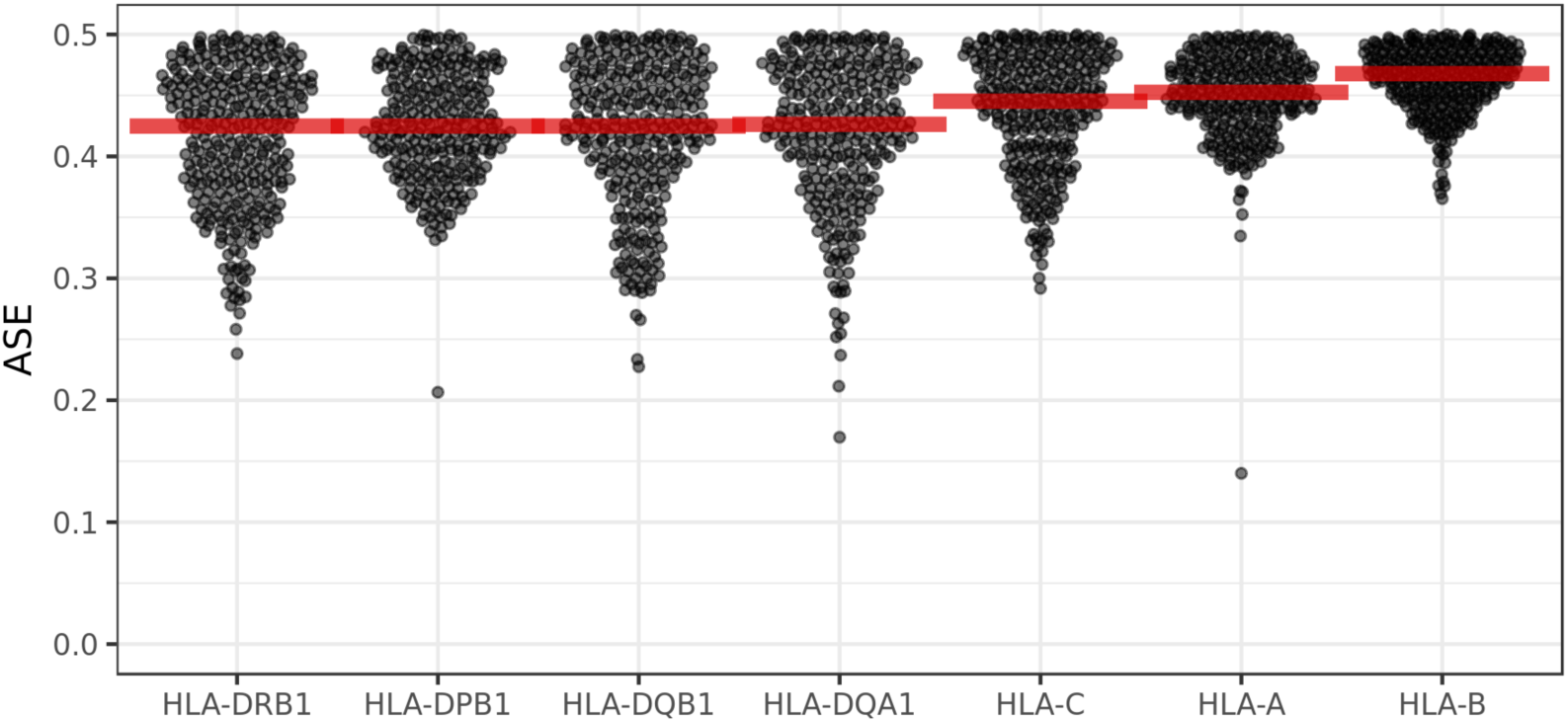
Allele-specific expression. ASE was measured as the proportion of the locus-level expression attributed to the less expressed HLA allele in heterozygous genotypes. Red horizontal bars indicate the median.

### Co-expression and haplotypic coordination in expression

Enhanced coordination of gene expression has been proposed as an advantage for gene clustering, as seen at the HLA region [1, 2]. We found a high correlation of expression both within the group of Class I and Class II genes, and lower levels between Class I and Class II genes, which are at least 1.2Mb apart (Figure 8A) [but see 19, for a result of no co-expression among Class I genes].

**Figure 8:**
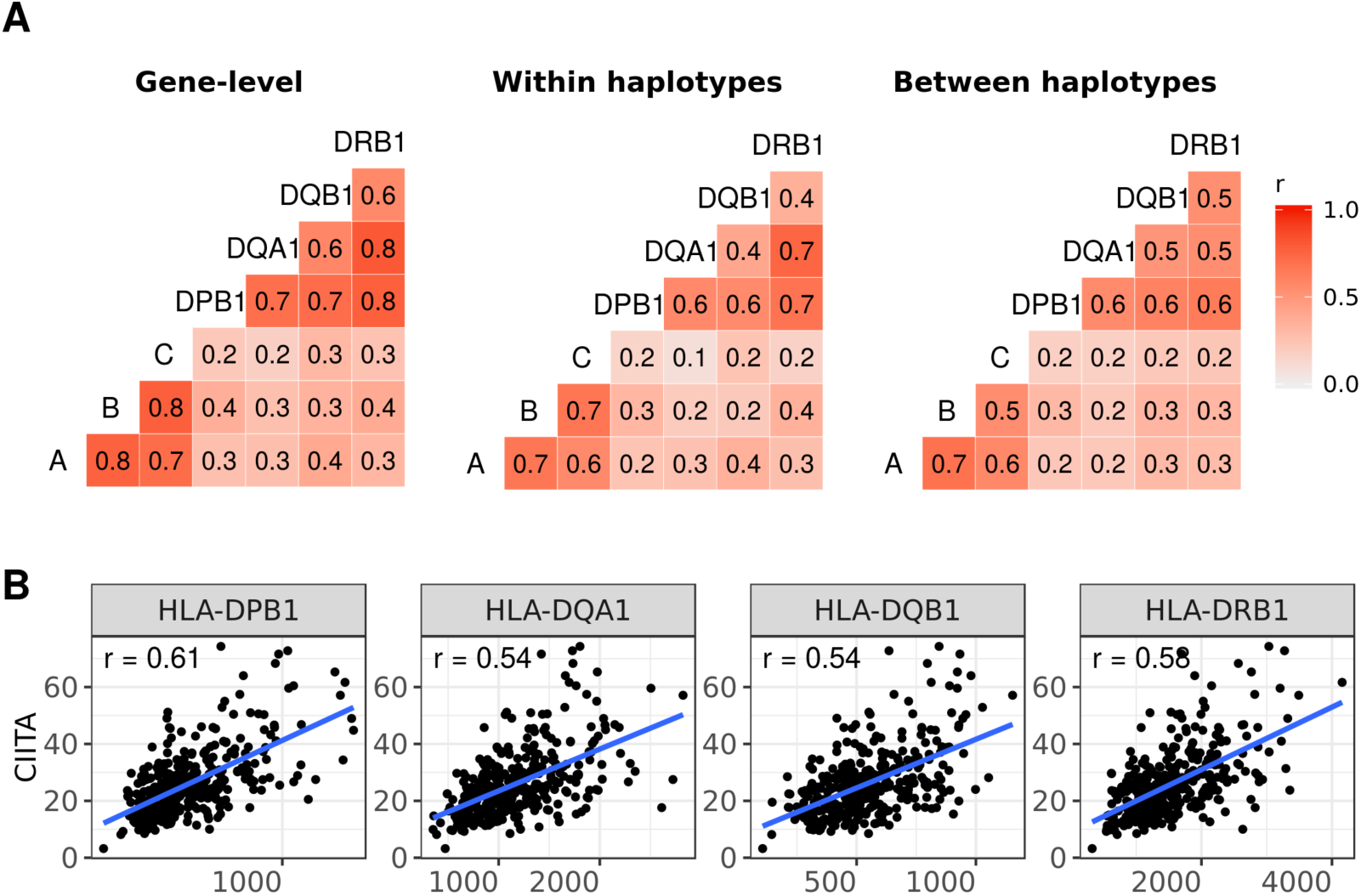
Co-expression patterns for HLA genes. **A** Pairwise correlation of expression estimates at the gene-level, for alleles from within the same haplotype, and for alleles on different haplotypes. **B** An example of a factor that contributes to correlation of expression at the gene level. *CIITA* is located on chromosome 16, and its protein regulates the expression of Class II genes.

A possible cause for co-expression of genes which are physically close to each other is that regulatory activity is structured in domains (CRDs, for Cis Regulatory Domains) [45]. Such domains comprise contiguous regions along a chromosome, and their existence predicts co-expression along haplotypes for genes associated with the same CRD. In fact, a previous study did find evidence that expression of some genes in the MHC region was a feature associated with haplotype membership [20].

We used our inferences of HLA haplotype structure to investigate if there is an haplotypic effect on coordination of expression among nearby HLA genes. Specifically, we tested the hypothesis that co-expression is stronger between alleles located within the same haplotype than between those on different haplotypes. We did not find a consistently higher correlation of expression for alleles on the same haplotype (correlation within haplotypes being higher in only 4 out of 9 locus pairs surveyed within Class I or Class II, Figure 8A).

This result suggests that correlation of expression among HLA loci in LCLs is a result of factors acting at the gene level, and is not driven predominantly by properties of the haplotype. For example, we observe correlation of expression between Class II genes and *CIITA*, the Class II Major Histocompatibility Complex Transactivator. This correlation is not driven by proximity *(CIITA* is on chromosome 16), but rather by a *trans* regulatory mechanism (Figure 8B).

## Discussion

The contribution of HLA to normal and disease phenotypes goes beyond peptide specificity, and includes other factors which influence the strength of immune responses, such as HLA expression levels. However, we are still only starting to understand the regulation of expression of these genes, a consequence of their extreme polymorphism which imposes challenges to the methodologies available to measure mRNA and protein levels.

There has been an increasing effort to adapt existing methods used to analyze RNA-seq data so as to allow reliable *in-silico* HLA typing ([55] and other methods reviewed in [56]), and accurate expression estimates for HLA alleles and genes [30, 33] in large datasets.

In this paper, we present HLApers, an HLA-personalized pipeline that reliably quantifies HLA expression based on RNA-seq. Our pipeline differs from previous approaches used to quantify HLA expression by incorporating a Maximum-Likelihood estimator to deal with instances of reads mapping to multiple alleles or loci, following a strategy that has been widely used in RNA-seq studies [26, 57].

Our pipeline is implemented in a way that allows different alignment strategies to be used. We show that the accuracy of expression depends on the sequences contained in the index, and is less dependent on the specific implementation used. We use simulations and data analyses to evaluate the performance of the HLApers pipeline, and explore how the use of this pipeline influences downstream analyses.

The impact of using an HLA-personalized index on expression estimates varies markedly among loci. We found that for *HLA-DQA1* there is a large difference between gene-level expression estimates obtained using the reference transcriptome and the HLA-personalized index. However, this difference is quite low for Class I genes, and intermediate for the other Class II loci (Figures 2 and 4). However, even for the class I genes, using the personalized index results in changes in expression estimates. Therefore, we asked whether the differences in expression obtained using the HLA-personalized pipeline have an impact on downstream analyses, such as eQTL mapping.

In the case of *HLA-DRB1*, the eQTLs identified using the HLA-personalized pipeline are markedly different from those obtained using the reference transcriptome. For the other loci we studied, although the eQTLs differ depending on whether the HLA-personalized index is used, the same biological signal was being captured (Table S2). The p-values and the causality of the eQTLs obtained with the personalized pipeline are only modestly better than when the reference transcriptome is used.

We provide a set of variants associated with independent effects on HLA regulation, and although it remains difficult to pinpoint the exact causal variants with a limited sample size, our analysis framework will be applicable to larger RNA-seq datasets.

Importantly, the HLA-personalized approach provides expression estimates at the HLA allele level, which is not a product of standard RNA-seq pipelines. We integrate allele level information with the eQTLs mapped for the genes, showing that the HLA allele is a relevant layer of information to understand the regulation of gene expression, because in some instances the regulatory architecture is linked to specific HLA alleles. This joint mapping of regulatory variants and assessment of expression of HLA alleles can illuminate the understanding of the HLA regulation, and contribute to disentangle specific contributions to disease phenotypes.

## Materials and methods

### Index supplemented with the HLA diversity

In order to create the index, we downloaded 16,187 nucleotide sequences for 22 HLA loci *(HLA-A, HLA-B, HLA-C, HLA-E, HLA-F, HLA-G, HLA-DMA, HLA-DMB, HLA-DOA, HLA-DOB, HLA-DPA1, HLA-DPB1, HLA-DQA1, HLA-DQB1, HLA-DRA, HLA-DRB1, HLA-DRB2, HLA-DRB3, HLA-DRB4, HLA-DRB5, HLA-DRB7, HLA-DRB8*) from the International Immunogenetics/HLA database *(IMGT*) (release 3.31.0 available at https://github.com/ANHIG/IMGTHLA).

For many alleles, sequence data is not available for the entire coding region (e.g., only for exons 2 and 3 for class I, and exon 2 for class II genes, which are called ARS exons). Because the lack of sequences for much of the coding region would cause the exclusion of many reads, including those mapping to the boundaries of the available exons, for each allele with partial sequence we used the available sequence to find the closest allele which has the complete sequence, and attributed the sequence from this allele. This is expected to introduce little bias both in either the genotyping step (because ARS exons are the most polymorphic and sufficient to distinguish specific alleles) or the expression estimation (because the non-ARS sequence attributed is likely very similar to the real one).

For the final index file, we replaced the HLA transcripts in the reference transcriptome (Gencode v25, primary assembly) with the HLA diversity described above. STAR’s module `genomeGenerate`, Salmon’s `index`, and kallisto’s `index` compile an index from these sequences.

### *In-silico* HLA typing

In order to select the alleles to be used in the HLA-personalized index, we observed that a simple procedure of selecting the 2 alleles with the largest number of estimated read counts after applying a zygosity threshold is sufficient to produce calls with accuracy of >95%. However, in order to avoid false homozygotes and false heterozygotes, we implemented additional steps.

First, we selected the top 5 alleles and applied an intra-lineage threshold of 0.25, meaning that only alleles which had at least 25% of the total expression in their lineage were considered for further steps. For each individual, we compiled an index containing only these (up to) 5 alleles and estimate their expression. We then determined if the individual was heterozygote at the lineage level by applying a threshold of 0.15 on lineage expression levels. The lead allele from each lineage was selected to compose the genotype. A zygosity threshold of 0.15 was applied to decide whether the genotype was heterozygous at the allele-level. For each locus, the reads mapped to the lead allele were removed, and another step of alignment and quantification was performed in order to determine if the second allele was real, or just noise due to extensive similarity to the lead allele. If the second allele had at least 1% of the locus read counts, it was kept, otherwise the genotype was considered to be homozygous for the lead allele.

The thresholds described above were chosen because they maximized the concordance with the Sanger sequencing typings [6], while also minimizing the rate of false homozygotes and heterozygotes.

### Expression quantification

We implemented two versions of the HLApers pipeline: (1) one using STAR (v2.5.3a) [34] to map reads followed by Salmon (v0.8.2)[36] to quantify the expression, and (2) using kallisto (v0.43.1)[35], which performs pseudoalignment and quantifications.

The quantification pipeline is structured in a two-stage process, first identifying the most expressed allele(s) at each HLA locus in order to infer the genotype which is present, and next quantifying expression for these inferred genotypes as well as for the rest of the transcriptome (Figure 1).

Reads were aligned directly to the transcriptome. STAR alignments were passed to Salmon for quantification (module `quant` under alignment mode), whereas kallisto directly produces quantifications with the `quant` module.

In both the HLA typing step (for which the index contains all HLA sequences in the IMGT database) and the quantification step (for which an HLA-personalized index is used), short reads can map to more than one locus, or more commonly to multiple alleles of the same locus (multimaps). The quantification methods we are using deal with multimaps by inferring maximum likelihoods estimates optimized by a expectation-maximization algorithm to probabilistically assign reads to each reference in the index, and also include models to account for sequencing bias.

For the mapping with STAR, we tuned parameters in order to avoid discarding multimaps and to accommodate mismatches. For quantification, we used all bias correction options available (`-seqBias` and `-gcBias` in Salmon, and `-bias` in kallisto).

### Simulation

We simulated transcriptome data for 50 randomly chosen GEUVADIS individuals. We used the Polyester package [41] and the read counts estimated for these samples to simulate RNA-seq experiments with library sizes of 30 million reads, sampling from a normal distribution of read start sites (with average fragment length of 250bp and sd of 25bp, and error rate of 0.005). Then we processed the simulated reads with STAR-Salmon to perform the quantifications using different indices:

1. HLA-personalized index (HLApers): Reference transcriptome with the annotated transcripts for HLA genes replaced with sequences from the personalized genotypes.
2. Reference transcriptome: Gencode v25 transcripts from the primary assembly of reference genome;
3. Reference Genome (GRCh38), considering only uniquely mapped reads.

To investigate the relationship between quantifications and sequence divergence with respect to the reference, we used the R function `adist` to calculate the proportion of mismatches between the HLA alleles carried by the individuals and the alleles in the reference genome.

### GEUVADIS reanalysis

We quantified HLA expression based on RNA-seq data for 358 European individuals, in samples of LCLs (Lymphoblastoid Cell Lines), as originally reported by the GEUVADIS Consortium [24] (we excluded samples from the original dataset which are not in the 1000 Genomes phase 3).

We performed expression quantification using the HLApers and reference transcriptome pipelines.

For the eQTL analysis, we used only autosomal genes which are expressed in a large proportion of samples, exploring the thresholds of *TPM >* 0 in at least 25%, 50%, or 75% of samples. In order to correct the expression data for technical effects, we sequentially removed the effect of the first 0 to 100 PCs and ran an eQTL analysis for each condition (Figure S5). The configuration of thresholds and number of PCs which maximized the eQTL discovery (at FDR = 5%) was considered. This resulted in the use of genes expressed in ≥ 50% of samples (19,613 genes), and 60 PCs.

The PCA analysis and data correction were performed with `QTLtools` v1.1 [43], using the modules `pca` and `correct` respectively.

For the genetic variant data, we used the 1000 Genomes Phase 3 biallelic variants, lifted to GRCh38 coordinates, after filtering for *MAF* ≥ 0.05 in the individuals included in this study (6,837,505 variants in total).

In order to control for population structure in the eQTL analysis, we ran a PCA on the variant genotype data and assessed the PCs which captured the structure. We used `QTLtools pca` requiring that variants should be at least 6kb apart. After visual inspection of the plots in Figure S6, PCs 1-3 were used as covariates in the eQTL analysis.

We used `QTLtools cis` to conduct the cis-eQTL analysis using the following model:

PCA-corrected and standard normal expression ~ SNPs + covariates (PCs for population stratification)

The permutation pass was performed with 1000 permutations and a cis-window of 1Mb. P-values were computed by beta approximation and significance was determined by running the script +`runFDR_cis.R` provided by `QTLtools` with FDR of 5%.

Multiple eQTLs with independent effects on a particular gene were mapped with a conditional analysis based on step-wise linear regression (see Supplementary method 8 in [43]). The method automatically learns the number of independent signals per gene and provides sets of candidate eQTLs per signal.

### Functional annotation of eQTLs

In order to investigate the putative function of the eQTLs we mapped, we investigated whether these eQTLs were present in ENCODE [47] regulatory elements annotated for lymphoblastoid cell lines (LCLs). We used three types of functional annotations: open chromatin regions given by DNAse footprinting, transcription factor binding sites (TFBS) assayed by ChlP-seq, and histone modifications.

### Regulatory Trait Concordance (RTC) analysis

We performed an RTC [44] analysis as described in [43] to investigate whether our eQTLs tagged the same causal variant as a GWAS variant or previously reported eQTL.

We downloaded the GWAS catalog data (v1.0.1) from https://www.ebi.ac.uk/gwas/api/search/downloads/alternative and selected associations with p-value < 10^−8^. We obtained the coordinates of recombination hotspots from http://jungle.unige.ch/QTLtools_examples/hotspots_b37_hg19.bed.

We applied the RTC module implemented in QTLtools (`QTLtools rtc`), selecting the HLA region only, using a D’ threshold of 0.5, and turning on the conditional flag (`-conditional`) to test all independent eQTLs for a gene.

### Haplotypic coordination of expression

To investigate whether there is a haplotypic coordination of expression at HLA, we used phased HLA genotype data to verify if there was more correlation of expression between alleles on the same haplotype than on different haplotypes.

We used PHASE [58] to determine the haplotype of each allele in the genotype, providing HLA allele designations and phased eQTL genotypes as input.

### Code availability

The HLApers pipeline is available at https://github.com/genevol-usp/HLApers. The entire analysis, including simulations, index compilation, quantification of expression, eQTL mapping, etc is available at https://github.com/genevol-usp/hlaexpression.

## Acknowledgements

Vitor Aguiar was supported by the São Paulo Research Foundation (FAPESP, fellowships #2014/12123-2 and #2016/24734-1). Jonatas Cesar was supported by the FAPESP fellowship #2015/19990-6.

